# CCR2^+^ neutrophils exhibit a proinflammatory phenotype and promote plaque destabilization

**DOI:** 10.1101/2025.08.06.669018

**Authors:** Merieme Farjia, Chang Pan, Quinte Braster, Patricia Lemnitzer, Nadja Sachs, Christian Schulz, Lars Maegdefesel, Oliver Soehnlein, Carlos Silvestre-Roig

## Abstract

**Background:** Atherosclerotic plaque destabilization is promoted by inflammatory cell recruitment, tissue cell death and mechanical weakening. Neutrophils are key instigators of vascular tissue injury and perpetuation of inflammation, and targeting their actions is a viable therapeutic opportunity. We here identify a distinct subset of activated neutrophils within atherosclerotic lesions that express the chemokine receptor CCR2, which directs their migration toward areas enriched with smooth muscle cells (SMCs) and contributes to plaque instability.

**Methods:** Flow cytometry and single cell transcriptomic analysis of CCR2^+^ neutrophils within murine and human atherosclerotic plaques. Tracking of CCR2^+^ neutrophil in hypercholesterolemic *Ldlr*^-/-^ mice and in mice reconstituted with *Ccr2*^*GFP/-*^ bone marrow. In vivo reactive oxygen species (ROS) and neutrophil extracellular trap (NET) analysis in lipopolysaccharide-induced peritonitis. In vitro migration assays of neutrophils deficient for or treated with specific antagonist against chemokine receptors. In vivo neutrophil recruitment and atherosclerotic plaque destabilization analysis upon CCR2 and CCL2 blockade in a model of advanced atherosclerosis in *Apoe*^*-/-*^ mice.

**Results:** CCR2^+^ neutrophils preferentially populate mouse and human atherosclerotic lesions and displayed a proinflammatory phenotype with enhanced capacity for ROS production and NET release. Genetic deletion or pharmacologic inhibition of CCR2 significantly reduces neutrophil migration and infiltration to the atherosclerotic lesion, reducing their presence in SMC-rich areas. Consistently, neutralization of the CCR2 ligand CCL2 decreased lesional neutrophil numbers and preserved fibrous cap integrity by increasing SMC content and decreasing overall plaque instability.

**Conclusions:** Our data suggest that a subset of CCR2-expressing neutrophils senses SMC-derived CCR2 ligands to infiltrate and destabilize atherosclerotic lesions. These results support the existence of neutrophil functional heterogeneity within the atherosclerotic lesions contributing to alterations of lesion stability. Specific targeting thereof may improve plaque stability without impacting host defense.

## Introduction

Neutrophils constitute the frontline of defense against infections but also incite tissue damage upon excessive inflammation. In advanced atherosclerosis, neutrophils continuously infiltrate the vascular tissue and cause tissue damage through the release of mediators such as reactive oxygen species (ROS), proteases or neutrophil extracellular traps (NETs), thus perpetuating the inflammatory response.^1^ This progressive noxious activity drives the progression of plaque development and increases its vulnerability. Mechanistically, we have previously shown that NETs released by lesional neutrophils contain extracellular histone H4, which can bind to smooth muscle cells (SMC) membranes and induce lytic cell death, leading to the weakening of atherosclerotic plaque.^2^ Interestingly, chemoattractants released by activated SMCs are responsible for neutrophil activation and NET release in the vicinity of SMCs^2^. Arterial migration of neutrophils during atherogenesis depend on different chemokine receptors such as CCR1, CCR5, CXCR2 or CCR2^3^. At the vascular site, CCR2 ligands trigger neutrophil activation with CCL7 initiating the release of NETs^2^; however, their role in neutrophil migration through the vascular tissue remains unknown. Notably, and despite CCR2 being classically expressed on inflammatory monocytes, emerging data indicate that neutrophils can upregulate CCR2 and migrate to cognate ligands under certain inflammatory conditions. Evidence of this functional expression of CCR2 in neutrophils was observed in mouse models and patient samples during sepsis ^4^ or rheumatoid arthritis^5^, being essential for the development of the disease. Given the growing recognition of neutrophil heterogeneity,^6^ it has yet to be determined whether specific neutrophil subpopulations within atherosclerotic plaques utilize the CCR2-CCL2 axis for recruitment and activation, thereby contributing to plaque instability.

Here, we hypothesize that a subset of CCR2 expressing neutrophils is recruited to atherosclerotic plaques via SMC-derived CCL2 signals, and that these neutrophils contribute to plaque inflammation and destabilization. We investigated this hypothesis by examining human carotid plaques and mouse models of atherosclerosis for the presence and phenotype of CCR2^+^ neutrophils. We further assessed the functional importance of the CCR2 axis in neutrophil recruitment, activation and plaque stability using transcriptional and functional, and intervention studies. Our findings reveal a distinct CCR2^+^ neutrophil subset in atherosclerosis and suggest a relevant neutrophil-SMC chemokine crosstalk that accelerates plaque vulnerability.

## Methods

### Ethics statement

All experiments complied with European animal-care guidelines and were approved by Regierung von Oberbayern and LANUV. The use of human blood and tissues was approved by the Ethical Committee of the TUM Klinikum rechts der Isar. Samples were coded and clinical information of patients pseudonymized.

### Mouse procedures

Sample sizes were determined by statistical power calculations. Female eight-to ten-week-old C57BL/6J mice were randomly assigned, with blinded data collection and analysis. Treatments and sacrifice were performed by cage with random order. Animals were housed according to guidelines with free access to food and water. Apolipoprotein E-deficient mice (*Apoe*^−/−^), Low density lipoprotein receptor (*Ldlr*^*-/-*^) were purchased from The Jackson Laboratory. *Ly6g*^*CRE*^ were intercrossed with *Apoe*^*−/−*^ and *Cxcr2*^*fl/fl*^. *Ly6g*^*CRE*^ *Cxcr2*^*fl/f*^, *Ccr1*^*-/-*^, *Ccr2*^*-/-*^, and *Cxcr4*^-/-^ were intercrossed with *Apoe*^*−/−*^. *Ldlr*^−/−^ recipient mice were lethally irradiated (2x 6.5 Gy) and reconstituted with bone marrow (BM) obtained from *Ccr2*^*GFP/-*^ mice to generate mouse chimaeras. In antagonist treatment, vulnerable atherosclerotic lesions were induced as described in^2^. In brief, mice were fed a high-fat diet (HFD; 21% fat and 0.15% cholesterol) for 11 weeks. Two weeks after the initiation of the HFD, a cast was placed around the left common carotid artery. CCR2 antagonist (RS504393, 5 mg/kg) was injected intravenously 2 hours before sacrifice to assess neutrophil recruitment. For CCL2 blockade, mice were fed with high-fat diet (HFD; 21% fat and 0.15% cholesterol) for 12 or 16 weeks. Subsequently, mice were treated daily with intraperitoneal injection of anti-CCL2 (20 µg per day, BioXcell, BE0185) or control IgG (20 µg per day, BioXcell, BE0091) during the last 4 weeks of the experiment to therapeutically treat pre-established atherosclerotic plaques. For LPS-induced peritonitis, mice were injected with 10 µg of LPS intraperitoneally for 24 hours.

### scRNA-seq analysis

Single cells suspensions from blood, BM and atherosclerotic tissue were counted with flow cytometry, normalized across conditions, tagged with hashtag antibodies specific for each organ. After extensive washing, cells were pooled, FACS-sorted as: DAPI^-^ CD115^+^ Ly6C^+^ for blood and BM, and for DAPI^-^ CD45^+^ aortic tissue. Sorted cells were loaded into the 10x Genomic platform and processed with 10X Chromium Next GEM Single-Cell 3′ Reagent Kit with Feature Barcoding technology and sequenced (Novogene). Raw reads were processed with cellranger 6.0.2 and aligned against Mus Musculus GRCh38 mm10.

Public datasets (GEO accessions GSE97310, GSE123587, GSE154817, GSE252243, GSE116240, GSM4981311) and own dataset were imported into R as a SingleCellExperiment object and combined after gene intersection, row metadata standardization, and cell annotation by experimental condition and organ of origin. QC metrics (total counts, detected genes and mitochondrial fraction) were computed using scater. Outliers (based on median absolute deviations), genes expressed < 6 cells or with average counts below 1×10^−5^ were filtered out. Data were normalized (library-size) and log-transformed (scran::logNormCounts), and high-variance genes (modelGeneVar) were calculated and used for principal component analysis (PCA). Batch effects were corrected using Harmony^7^ (10 top PCs), followed by Uniform Manifold Approximation and Projection (UMAP), shared nearest neighbor (SNN) graph calculation on the Harmony-corrected PCs and Louvain clustering. Cells were annotated using SingleR with the ImmGen reference and manual curation. Next, neutrophils were subset from the integrated object and normalization, PCA, Harmony correction, UMAP, and Louvain clustering was repeated. CCR2 expression was binarized (raw UMI ≥1) to define CCR2^+^ and CCR2− neutrophils.

Gene counts (genes in ≥ 10% of cells) were aggregated into a DGEList TMM-normalized (edgeR), and modeled with a quasi-likelihood NB fit (glmQLFit) using CCR2 status or cluster label. DEGs were identified with glmTreat (log_2_ FC ≥ 0.585, FDR < 0.05), ranked, and used to create gene-signature lists (CreateGeneSignatures). Gene-set enrichment analysis was performed via ESCAPE (UCell). Signature score differences between CCR2^+^ and CCR2− neutrophils were evaluated by Welch’s t-test with Benjamini–Hochberg correction (rstatix). Visualization plots were performed in ggplot2.

### Tissue processing

Mice were euthanized with a ketamine/xylazine overdose, followed by blood collection, perfusion (ice-cold PBS-EDTA, 5 mM) and organs harvesting for flow cytometry or immunostaining. Cells were washed, centrifuged, and resuspended in HANKs buffer (HBSS Mg^-^/Ca^-^+0.06% BSA+0.3 mM EDTA). BM was isolated by flushing the femur with 5 ml HANKs buffer using a 21G needle. Aortic arches and hearts were embedded unfixed in Tissue Tek O.C.T. compound. Human and mouse atherosclerotic plaques were digested in 1 ml RPMI with 10% FCS and 1.25 mg/ml Liberase at 37 °C for 1 hour.

### Flow cytometry

Red blood cells were lysed (150 mM NH_4_Cl, 10 mM KHCO_3_, 0.1 mM Na_2_EDTA) for 5 min at room temperature, and leukocytes were stained with antibodies to CD45 (30-F11), CD11b (M1/70), Ly6G (1A8), Ly6C (Hk1.4), CD115 (AFS98), Ly6B.2 (BIORAD, clone: 7/4), CCR2 (SA203G11), CD80 (16-10A1), CD11a (M17/4), CD86 (A17199A), CD14 (Sa14-2), CD74 (In1/CD74), CXCR2 (SA044G4), MHCII (M5/114) and C5ar (20/70) from Biolegend or BD Biosciences in staining buffer (20 min, 4 °C). Human blood was incubated with red blood cell lysis (5 min, room temperature + 5 min, 4°C) and cells were stained with antibodies for CD45 (2D1), CD11b (Ma/70), HLA-DR (L243), CD64 (10.1), CD14 (63D3), CD16 (3G8), CD3(HIT3a), CD66b (G10F5), CCR2 (K036C2) from BioLegend. Flow cytometry was performed using the LSR Fortessa (Beckton Dickinson) and Cytek Aurora (Cytek Biosciences) and data were analyzed using FlowJo software (Beckton Dickinson).

### Cell function analysis

24h post-LPS injection, cells from BM, blood and peritoneal lavage we isolated. For ROS production analysis, cells were incubated with DHR (Dihydrorhodamin 123, Invitrogen) for 30 min at 37°C. For NETosis analysis, cells were intracellularly stained with anti-citH3 primary antibody (Abcam, 1:300) and anti-rabbit secondary antibody (Sigma, 1:500).

### Histology and immunofluorescence

Brachiocephalic arteries (7 µm) cryosections were histologically stained with hematoxylin and eosin (H&E) in 40 µm intervals. Total collagen content was assessed by Pricrosirius Red staining in consecutive sections. For immunofluorescence staining, cryosections were fixed with cold acetone, blocked with 5% goat serum/phosphate buffered saline and, stained overnight at 4 °C with primary antibodies to rabbit anti-mouse CD68 (Abcam, 1:200), rat anti-mouse Ly6G (BD, 1:200), mouse anti-mouse smooth muscle actin (SMA)-FITC or -Cy3 conjugated (Sigma, 1:500), anti-mouse CCL2 (eBioscience, 1:100). After extensive washing, sections were incubated with secondary antibodies conjugated with Dylight 488, DyLight 550 or DyLight 650 (Thermo Fisher, 1:500) and counterstained to visualize nuclei using DAPI (Molecular Probes). Immunofluorescence sections were imaged using a Leica Thunder and TCS SP8 (Leica Microsystems). Histological sections were quantified by computer-assisted morphometric analysis using ImageJ software (National Institutes of Health).

### Murine plaque analysis

To assess plaque vulnerability, intima, media and necrotic core (NC) area was analyzed in H&E-stained sections. The NC was defined as the area devoid of nuclei underneath a formed fibrous cap. Collagen content and fibrous cap thickness were measured on Pricosirius Red-stained sections. Vulnerability Plaque Index (VPI) was calculated as VPI = (% NC area + % CD68 area) / (% SMA area + % collagen area).

### Cell culture and activation

Mouse vascular aorta/smooth muscle cells (MOVAS; ATCC CRL-2797) were cultured in DMEM (Gibco) with 10% fetal bovine serum, 0.2 mg/ml G418, and 5 mM sodium pyruvate. Cells were incubated at 37 °C with 5% CO_2_. For activation, MOVAS were treated with 10 ng/ml recombinant murine Platelet-derived growth factor (PDGF)-BB for 6 hours, then washed and placed in fresh medium for 24 hours. Supernatants were collected, centrifuged (300g, 5 min, 4 °C), and frozen.

### Neutrophil migration

Blood was collected from C57BL/6J mice, lysed for 5 min at room temperature and stained with Ly6G-PE (1A8, BioLegend) to label circulating neutrophils. Labelled cells were seeded (30 min) on a collagen-coated coverslip and mounted on a Zigmond glass slide chamber. A gradient of PDGF-BB activated SMC supernatants was created and images were acquired over 30 min in 30 s intervals using a climate chamber fluorescence microscope (20X dry objective, Leica, DMi8). For blocking of chemotactic receptors, inhibitors to CCR1 (BX471, 1uM), CCR2 (RS504393, 3.6 µM), CCR5 (DAPTA, 0.1 µM), CXCR2 (SB225002, 300 nM) or CXCR4 (AMD3100, 5 µg/mL) were added to supernatants. Speed and displacement were calculated as previously described^2^.

## Statistics

Statistical analyses were conducted with GraphPad Prism 10. Outliers (Q = 1%) were excluded using the ROUT function and data normality checked by the D’Agostino-Pearson test. A two-tailed unpaired or paired t-test (one variable) or one-way ANOVA with Tukey or Dunnett’s correction (>2 variables) was used, with p < 0.05 (95% CI). Results are expressed as mean ± SEM.

## Data and code availability

The data that support the findings of this study are available from the corresponding authors upon reasonable request.

## Declaration of generative AI and AI-assisted technologies

ChatGPT 4o was used during the generation and improvement of the R code. ChatGPT 4o and DeepL write was used to improve English grammar. After using this tool, the authors reviewed and edited the content as needed and take full responsibility for the content of the publication.

## Results

### CCR2^+^ neutrophils populate atherosclerotic lesions in humans and mice

We first investigated the presence of neutrophils expressing the CCR2 chemokine receptor within human blood and atherosclerotic plaques (**Figure 1A)**. Analysis of paired blood and plaque samples was performed in patients with cardiovascular disease (CVD) undergoing carotid endarterectomy, as well as in blood from healthy donors. In healthy individuals (n=7), circulating neutrophils (gated as CD45^+^CD11b^+^CD66b^+^CD16^+^ cells) showed a low frequency of CCR2^+^ neutrophils without a significant increase in CVD patients (**Figure 1B**). On the other hand, CCR2^+^ neutrophils were present in the atherosclerotic carotid in greater numbers (∼1.14–24.10%) compared to patient-matched blood samples (**Figure 1C**), indicating a specific accumulation or in situ generation of this subpopulation in atherosclerotic tissues. To corroborate these findings in an experimental model, we employed a reporter mouse expressing GFP under the control of *Ccr2* promoter (**Figure 1D**). *Ldlr*^-/-^ mice were transplanted with bone marrow (BM) from *Ccr2*^*GFP/-*^ donors followed by high-fat diet (HFD)-feeding for 12 weeks to develop advanced atherosclerosis. Flow cytometric analysis of BM, blood, and aorta revealed the presence of CCR2-GFP^+^ neutrophils in all compartments (**Figure 1E/F**). As observed in humans, only a small fraction of neutrophils in blood (∼2.80%) and BM (∼5.81%), whereas a significantly larger fraction of aortic neutrophils was CCR2-GFP^+^ (∼14,51%, **Figure 1E/F**), confirming an enrichment of a subset of neutrophils in the vascular atherosclerotic tissue expressing the non-canonical chemokine receptor CCR2.

**Figure 1.**
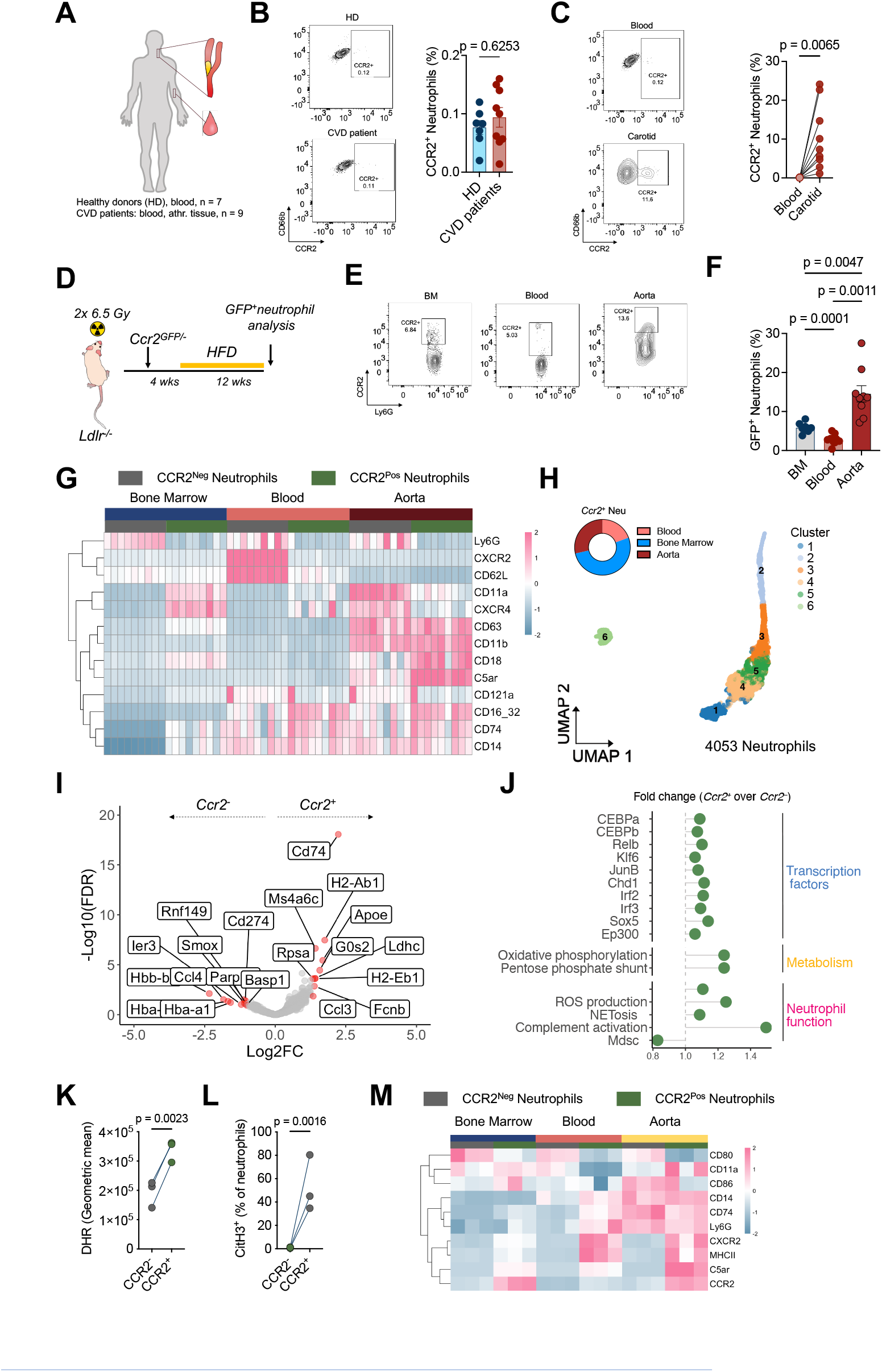
CCR2^+^ hyperactive neutrophils populate mouse and human atherosclerotic lesions. **A-C**. Analysis of CCR2^+^ neutrophils in blood and atherosclerotic lesion from healthy donors (HD) and patients with cardiovascular disease (CVD). **A**. Experimental design. Blood was drawn from HD (n=7), and CVD patients (n=9). One HD and CVD were analysed at each time. Atherosclerotic tissue was processed from carotid endarterectomies. **B**. Flow cytometry analysis of blood CCR2^+^ neutrophils (CD45^+^CD11B^+^CD66B^+^CD16^+^). Unpaired student t-test. **C**. Paired flow cytometry analysis of blood and atherosclerotic tissue. Paired student t-test. **D-F**. Analysis of CCR2-GFP^+^ neutrophils in hypercholesterolemic mice. **D-G**. *Ldlr*^*-/-*^ mice (n=9, 1 independent experiment) were lethally irradiated and reconstituted with bone marrow (BM) cells from CCR2^GFP/-^ mice. 4 weeks after reconstitution, mice were fed with HFD for 12 weeks. **E/F**. Flow cytometry analysis of CCR2-GFP^+^ neutrophils in BM, blood and aorta. **E**. Representative plot. **F**. Percentage of GFP^+^ neutrophils in indicated organs. **G**. Heatmap showing scaled mean fluorescence intensity from indicated cell surface markers in BM, blood and aorta from CCR2-GFP^+^ or CCR2-GFP^-^ neutrophils. **H**. Uniform Manifold Approximation and Projection (UMAP) of aggregated neutrophils from BM, blood and atherosclerotic tissue. Pie chart displays proportion of *Ccr2*^+^ neutrophils within these organs. **I**. Volcano plot showing significantly expressed genes (red dots) between *Ccr2*^-^ and *Ccr2*^+^ neutrophils. **J**. Lollipop plot showing fold-change of indicated transcriptomic scores in *Ccr2*^+^ over *Ccr2*^-^ neutrophils (adjusted p-value <0.05). **K-M**. C57BL/6J mice (n=3, 1 independent experiment) were injected with 10 µg of LPS i.p for 24 h. **K**. Flow cytometry analysis of ROS production in CCR2^+^ and CCR2^-^ peritoneal neutrophils. Paired student t-test. **L**. Flow cytometry analysis of percentage of citH3^+^ within CCR2^+^ and CCR2^-^ peritoneal neutrophils. Paired student t-test. **M**. Heatmap showing scaled mean fluorescence intensity from indicated cell surface markers in BM, blood and peritoneum from CCR2^+^ and CCR2^-^ neutrophils. Data are shown as mean±SEM.

### CCR2 ^**+**^ **neutrophils exhibit a pro-inflammatory phenotype**

To profile CCR2-GFP^+^ neutrophils, we investigated the surface receptor expression of this subpopulation respective to CCR2^-^ neutrophils in the BM, blood, and aorta of atherosclerotic mice. Using a broad panel of activation and maturation markers, we observed that CCR2-GFP^+^ neutrophils were phenotypically distinct showing reduced CD62L and CXCR2 expression, indicating an activated phenotype and reduced maturation. However, in the atherosclerotic lesion, CCR2-GFP^+^ neutrophils expressed elevated levels of CD18 and a tendency for increased Mac-1 (CD11b) integrins suggesting increased adhesive capacity (**Figure 1G**). Interestingly, complement 5 receptor (C5ar) was also highly expressed in CCR2-GFP^+^ neutrophils supporting an enhanced activation and recruitment capacity. These data suggest that neutrophils expressing CCR2 represent an activated and adherent subpopulation within the core of the atherosclerotic lesion.

We then performed a single-cell RNA sequencing (scRNA-seq) analysis of BM, blood and atherosclerotic arteries from *Apoe*^*-/-*^ fed with high-fat-diet for 16 weeks to further define this CCR2^+^ neutrophil subset. In addition, we combined these data set with existing datasets from atherosclerotic lesions (see methods). After quality control and data integration using Harmony package^7^, we performed cell annotation and focused on neutrophils (**Figure 1H**). Unsupervised clustering showed 6 different clusters based on maturation gene expression. We then selected cells expressing RNA transcripts for *Ccr2* (**Figure 1H**) and performed a pairwise analysis comparing *Ccr2*^*+*^ versus *Ccr2*^*-*^ neutrophils (**Figure 1I**). Analysis of differential gene expression showed an increased expression of genes associated to inflammation such as *Ccl3* or *Fcnb*, as well as a prominent phenotype associated with antigen presentation (*Cd74, H2-Ab1, H2-Eb1*) in CCR2+ neutrophils. These surprising results are in accordance with the increased expression of CD74 at protein level (**Figure 1G**). To gain insights into the functional phenotype of this subpopulation, we analyzed transcriptional scores associated with transcription factor targets, metabolism and neutrophil function (**Figure 1J**). *Ccr2*^*+*^ neutrophils displayed an increased expression of targets for known transcription factors involved in neutrophil activation and effector function such as RELb, JUNb or IRF1 and IRF2. Moreover, this subpopulation showed increased oxidative phosphorylation and pentose phosphate pathway scores, which is linked to increased ROS production and NETosis (**Figure 1J**). Finally, transcriptional score for complement activation showed the higher score and confirmed observed increased levels of C5ar (**Figure 1G**). These transcriptomic signatures reinforce the concept that *Ccr2*^+^ neutrophils are an inflammatory subset that may actively contribute to lesion pathology. Finally, we assessed the ability of this subpopulation to generate inflammatory mediators such as ROS and NETs (**Figure 1K/L**). To obtain sufficient neutrophil numbers, we employed a mouse model of LPS-induced peritonitis. Consistent with our phenotypic and transcriptomic profiling, recruited CCR2^+^ neutrophils to the peritoneum expressed elevated levels of ROS (**Figure 1K**) and NETs (**Figure 1L**). Furthermore, this subpopulation expressed higher levels of antigen-presenting markers CD74 and MHC-II, and complement receptor C5ar (**Figure 1M**), thus confirming this phenotype across inflammatory models.

### Neutrophil recruitment to plaques depends on the CCL2-CCR2 chemokine axis

The presence of lesional CCR2^+^ neutrophils prompted us to test whether CCR2 is functionally required for neutrophil chemotaxis within the atherosclerotic lesions. In vitro migration analysis showed a striking impaired migration of neutrophils deficient for CCR2 compared to controls, with reduced velocities and migration distances (**Figure 2A-C**). This reduction was also found in absence of the canonical receptors CXCR2 and CCR5, whereas no effect was found in CCR1 and CXCR4 deficient cells. We next treated neutrophils with specific receptor antagonists during the migration assay to pharmacologically validate these findings. Consistent with the genetic data, blockade of CCR2 markedly reduced neutrophil migration efficiency (**Figure 2D–F**). Inhibition of other receptors, including CCR5, CXCR2, CXCR4 but not CCR1 also reduced neutrophil migration. In an in vivo setting, blockade of CCR2 led to a significant reduction in the number of lesional neutrophils in a model of advanced atherosclerosis (**Figure 2G**), supporting the importance of this receptor in neutrophil arterial migration. Interestingly, this blockade also increased the distance to SMC-rich areas (**Figure 2I**) and reduced the presence of neutrophils in these areas (**Figure 2J**), in line with our previous results showing that SMC-derived chemokine gradients recruit neutrophils into close proximity^2^. At the atherosclerotic lesion, immunostaining analysis of the ligand CCL2 showed a strong expression within endothelial cells and transitioning SMCs (SMA^+^CD68^+^), suggesting that these cells are important in the recruitment and positioning of neutrophils within the atherosclerotic lesion (**Figure 2K**).

**Figure 2.**
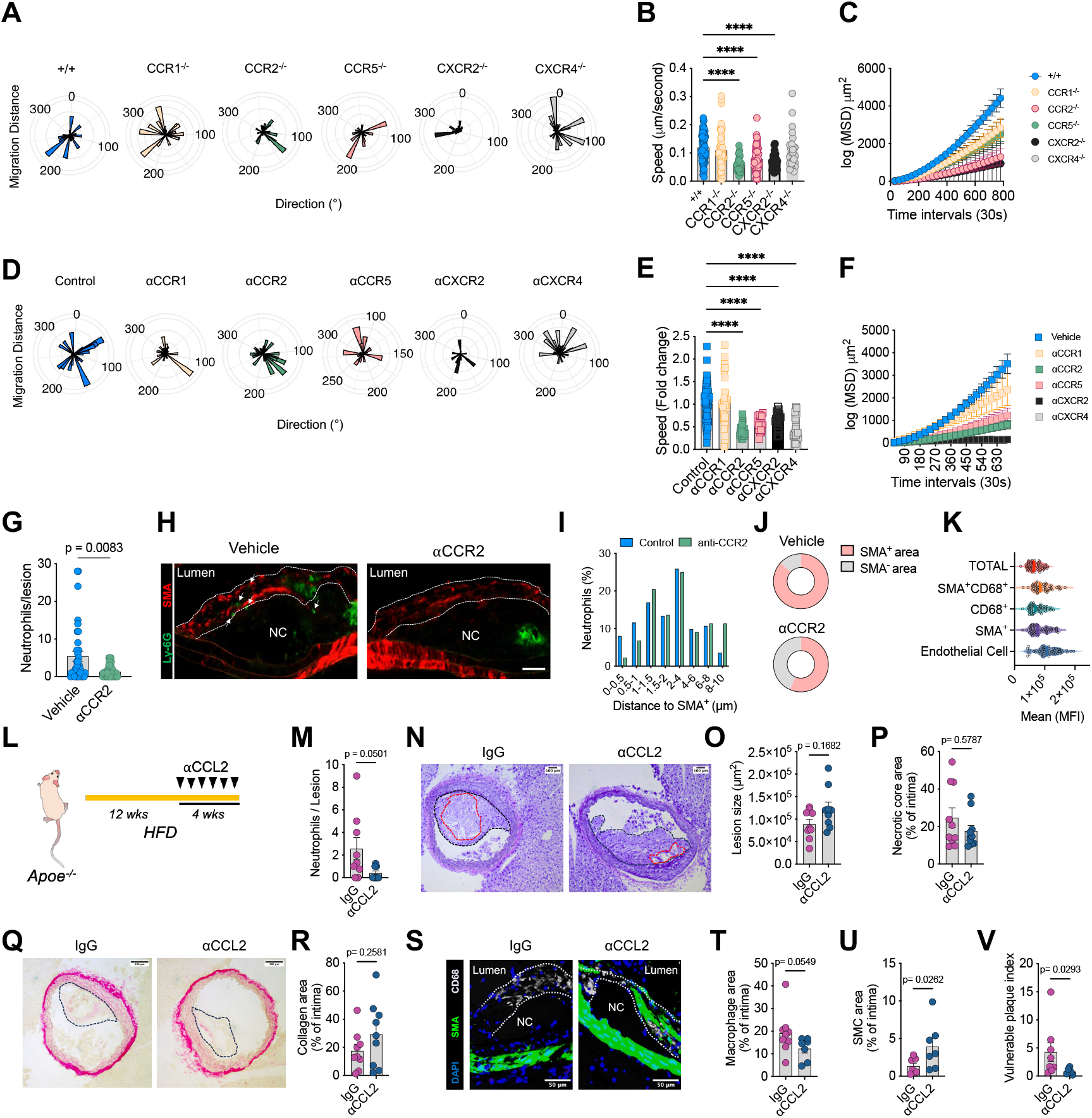
CCR2-CCL2 axis regulates neutrophil arterial migration and atherosclerotic plaque vulnerability. **A-C**. Neutrophil migration of cells deficient for CCR1, CCR2, CCR5, CXCR2 and CXCR4 (n=30-89 cells, 1-2 independent experiment/group). **A**. Rose plots showing migration distance of indicated genotypes. **B**. Migration speed (**B**) and Mean Square Displacement (MSD, **C**) analysis of neutrophils of indicated genotypes. **D-E**. Neutrophil migration of cells in presence of specific antagonists for CCR1, CCR2, CCR5, CXCR2 and CXCR4 (n=19-110 cells, 1-2 independent experiment/group). **D**. Rose plots showing migration distance of indicated treatments. Migration speed (**E**) and Mean Square Displacement (MSD, **F**) analysis of neutrophils of indicated treatments. **G-J**. *Apoe*^*-/-*^ mice (n=5-6/group, 1 independent experiment) were fed with high-fat diet (HFD) for 11 weeks. Advanced lesions were generated by insertion of a shear stress modifier around the carotid artery. **G**. Quantification of neutrophils by confocal immunofluorescence (Ly-6G^+^). **H**. Representative immunofluorescence micrograph showing lesional neutrophils (Ly-6G^+^, green) and SMCs (Smooth muscle actin (SMA)^+^, red). Scale bar, 50 µm. **I**. Percentage of neutrophils within binned distances towards nearest SMA^+^ smooth muscle cell (SMC). **J**. Neutrophil distribution in SMA^+^ and SMA^-^ SMC area. **K**. Mean fluorescence intensity of CCL2 expressed by the indicated cell types in the lesion. **L-V**. *Apoe*^*-/-*^ mice (n=9/group, 1 independent experiment) were fed with high-fat diet (HFD) for 11 weeks. During the last 4 weeks, mice were injected daily with anti-CCL2 (20 μg) or IgG as control. **L**. Experimental design. **M**. Quantification of neutrophils by immunofluorescence (Ly-6G^+^). **N**. Representative picture of HE staining of indicated treatments and quantification of lesion size. **O**. Necrotic core size. **P**. Representative picture of Pricrosirius Red staining and quantification of collagen area. **Q**. Representative confocal micrograph showing macrophages (CD68^+^, green) and SMC (SMA^+^, red). Quantification of macrophage (**R**) and SMCs (**S**) and vulnerable plaque index (**V**).

### Blocking CCL2 in vivo reduces neutrophil infiltration and improves plaque stability

Finally, we tested whether interrupting the CCR2-CCL2 axis in vivo would modulate neutrophil accumulation and plaque vulnerability. *Apoe*^*-/-*^ mice were fed with HFD for 16 weeks and then treated during the last 4 weeks with either a neutralizing antibody against CCL2 or an isotype control IgG (**Figure 2L**). Anti-CCL2 therapy led to a reduction in lesional neutrophils compared to controls, as quantified by Ly-6G^+^ immunostaining (**Figure 2M**). We next evaluated plaque composition and stability in treated and control mice. Although overall lesion size showed a modest decreasing trend with CCL2 blockade (**Figure 2N**), the most pronounced effects were on markers of plaque stability. anti-CCL2–treated plaques tended to reduce necrotic core area (**Figure 2O**) and increased collagen content (**Figure 2P**) relative to controls. Concomitantly, we observed a significant increase in SMC content and a non-significant reduction in macrophage content upon CCL2 blockade (**Figure 2Q-S**). To integrate these features, we calculated a vulnerable plaque index showing a substantial reduction upon blocking CCL2 (**Figure 2T**), indicating improved plaque stability.

## Discussion

Neutrophils infiltrate advanced atherosclerotic lesion and instigate cell death, thus driving plaque instability. We, here, identify a neutrophil-driven mechanism of inflammation in atherosclerosis, centered on a CCR2^+^ neutrophil subset that infiltrates plaques and promotes destabilization. We show that both mice and humans with advanced atherosclerosis contain neutrophils expressing the receptor CCR2 with preferential location in SMC-rich areas. These CCR2^+^ neutrophil subset are endowed with an activated phenotype and depend on the CCR2-CCL2 axis to infiltrate the atherosclerotic lesion and drive plaque instability.

Our previous work demonstrated a neutrophil-SMC detrimental interaction, where activated SMCs through the release of CCL7 prime neutrophils for NET release^2^. Interestingly, we here find that this attraction depends on CCL2 release and requires functional CCR2 expression in neutrophils. This chemotactic axis, despite being canonical for monocyte-macrophage migration, can also regulate neutrophil migration under inflammatory conditions^3, 5, 8^. Although neutrophils primarily rely on other receptors such as CXCR2 to reach the inflammatory site, the variable expression of chemokine receptors might favor selective recruitment of specific neutrophil subpopulations. In line with this idea, we identify that a subpopulation of CCR2^+^ neutrophil is enriched SMC-rich areas from mouse and human atherosclerotic lesions. This population is characterized by the surface expression of adhesion molecules such as CD18 and the complement receptor C5ar, which can cause the arrest ICAM1 or ICAM2-expressing SMCs. Interestingly, C5a is known to induce neutrophil arrest via beta2 integrins into the inflamed joint^9^, a mechanism that can similarly facilitate neutrophil-SMC interaction in the atherosclerotic lesion. Accordingly, our transcriptomic profile of lesional neutrophils supports this complement activation signature that aligns with an increased ROS and NETosis phenotype observed in neutrophils interacting with SMCs^2^, suggesting a mechanistic link between both processes. In addition to this pro-inflammatory phenotype, CCR2^+^ neutrophil express antigen-presenting surface proteins such as CD74 or transcripts such as *Cd74, H2-Ab1, H2-Eb1*, suggesting a potential capacity to interact with the adaptive immune system to promote atherosclerosis^10^. This specific phenotype was confirmed in a LPS-induced peritonitis model, where CCR2^+^ neutrophils overexpressed MHC-II and C5ar. Interestingly, this activated subpopulation exhibited an elevated capacity for ROS production and NETs release, thus confirming our transcriptomic data and supporting the pro-destabilizing properties of this subpopulation.

The concept of CCR2^+^ neutrophils adds to the evolving understanding of neutrophil heterogeneity in cardiovascular disease^11^, which remains understudied. Traditionally, neutrophils were thought to lack CCR2, however, inflammation can alter neutrophil receptor profiles in sepsis^4^, tumor^12^, or arthritis^5^ where neutrophil CCR2 expression has been evidenced. Our study first implicates CCR2^+^ neutrophils in atherosclerosis. CCR2^+^ neutrophils populate human and mouse atherosclerosis plaques suggesting that this subset expands or is preferentially recruited during chronic arterial inflammation. Hence, during advanced plaque development, neutrophil recruitment may be governed by tissue-derived CCR2 ligands gradients produced by intimal cells to attracting them towards specific regions such as the SMC-rich fibrous cap. Recently, SMC-derived CCL2 re-organizes macrophages around the fibrous cap protecting fibrous cap integrity^13^. Interestingly, and contrasting to our data, intravenous administration of blocking antibodies against CCL2 reduced SMC (ACTA2^+^) content without affecting any other parameter. Differences in the route or frequency of administration as well as duration of the model of advanced atherosclerosis might explain these differences. Thus, therapies targeting the CCR2 axis could have dual effects by affecting both macrophage survival and neutrophil contributions to plaque destabilization. A shift towards increased neutrophil recruitment respect to monocytes and reprogramming towards CCR2-expressing subsets may result in increased plaque destabilization. Considering this last possibility, in situ priming by local cues may drive the acquisition of this phenotype, a process that might depend on TLR4 signaling as found to be induced by LPS^4^ or to be absent in TLR4-deficient mice^14^. Whether atherosclerotic molecules signal through this pathway to drive the generation of CCR2^+^ neutrophils requires further investigation.

Altogether, this study provides evidence of a subset of CCR2^+^ neutrophils populating atherosclerotic plaques and guided by CCR2 chemotactic ligands released by SMCs, where they exacerbate inflammation and plaque instability. Targeting CCR2^+^ neutrophils neutrophil recruitment or reprogramming represents a promising strategy to protect the fibrous cap and prevent plaque destabilization.

## Acknowledgments

We thank Matthias Guntzer for (Institute for Experimental Immunology and Imaging, University Hospital, University Duisburg-Essen) for providing the *Ly6g*^*-/-*^ mouse model.

